# Genome-wide association identifies candidate genes for drought tolerance in coast redwood and giant sequoia

**DOI:** 10.1101/2021.10.25.465813

**Authors:** Amanda R. De La Torre, Manoj K. Sekhwal, Daniela Puiu, Steven L. Salzberg, Alison Dawn Scott, Brian Allen, David B. Neale, Alana R.O. Chin, Thomas N. Buckley

**Author notes:** Corresponding authors: Amanda De La Torre,; David B. Neale. Shared first authorship (Amanda R. De La Torre, Thomas N. Buckley). Department of Chromosome Biology, Max Planck Institute for Plant Breeding Research, Cologne, NRW, Germany.

## Abstract

Drought is a major limitation for survival and growth in plants. With more frequent and severe drought episodes occurring due to climate change, it is imperative to understand the genomic and physiological basis of drought tolerance to be able to predict how species will respond in the future. In this study, univariate and multitrait multivariate GWAS methods were used to identify candidate genes in two iconic and ecosystem-dominating species of the western US – coast redwood and giant sequoia – using ten drought-related physiological and anatomical traits and genome-wide sequence-capture SNPs. Population level phenotypic variation was found in carbon isotope discrimination, osmotic pressure at full turgor, xylem hydraulic diameter and total area of transporting fibers in both species. Our study identified new 78 new marker × trait associations in coast redwood and six in giant sequoia, with genes involved in a range of metabolic, stress and signaling pathways, among other functions. This study contributes to a better understanding of the genomic basis of drought tolerance in long-generation conifers and helps guide current and future conservation efforts in the species.

**Significance Statement:** Climate change brings more frequent and severe drought events that challenge the survival of natural populations of plants. While most of our knowledge about drought tolerance comes from annual and domesticated plants, the genomic basis of drought tolerance in long-generation trees is poorly understood. Here, we aim to fill this gap by identifying candidate genes in two conifer species, coast redwood and giant sequoia.

## INTRODUCTION

Understanding the genomic basis of phenotypic trait variation and its distribution across a species’ range is indispensable to predict species response to global climate change and to develop conservation and management guidelines (Bellard *et al*., 2012; Razgour *et al*., 2019). This has become an urgent need in the western United States, where longer and more severe drought events have resulted in massive tree mortality in the last ten years (Allen *et al*., 2010; Hicke *et al*., 2015; Adams *et al*., 2017; Stephenson *et al*., 2018; Fettig *et al*., 2019). Drought stress, manifesting as low soil water content and/or high evaporative demand, poses significant challenges to the establishment, development, growth and survival of long-generation tree species such as conifers (Adams and Kolb, 2005), and also predisposes trees to pathogens and pests (Jactel *et al*., 2012; Gaylord *et al*., 2013). Despite the economic importance of conifers and their dominance in global arid, semi-arid, montane and circumpolar zones, the genomics of drought and thermal tolerance have received little attention and lag behind studies in other plant species (Moran *et al*., 2017).

Conifer species have large genome sizes (8-34 Gb; Murray *et al*., 2004; De La Torre *et al*., 2014), and large genetic-to-physical distance ratio (>3000 kb/cM). Linkage disequilibrium (LD) in coding regions rapidly decays within a short distance, which complicates the identification of genes responsible for phenotypic variation (Neale and Savolainen, 2004). Another challenge of genotyping many individuals is the need to use a massive number of genome-wide markers in large-genome trees such as conifers. The development of high-throughput systems such as next generation sequencing (NGS) and SNP arrays should help overcome this difficulty, since they allow rapid and cost-effective genotyping over a massive number of SNPs (McCarthy *et al*., 2008). In addition, the rapid advancement of genome sequencing and bioinformatics approaches have opened the door to more comprehensive assessments of population-level diversity (McGuire *et al*., 2020). Association mapping principally exploits evolutionary recombination at the natural population level (Myles *et al*., 2009). A mixed linear model (MLM) method (Yu *et al*., 2006) was proposed to better control for population structure and the imbalanced kinships among various individuals (Pritchard *et al*., 2000). Until recently, determining the molecular basis of heritable trait variation has been challenging in conifer species, and genome-wide association studies have been limited to a few species and traits (Lu *et al*., 2017; Baison *et al*., 2020; Elfstrand *et al*., 2020; Weiss *et al*., 2020; Chen *et al*., 2021; De La Torre *et al*., 2021a). For example, association studies of drought tolerance have only been performed with pre-selected candidate genes (Gonzalez-Martinez *et al*., 2008; Eckert *et al*., 2010; Cumbie *et al*., 2011; Trujillo-Moya *et al*., 2018; Depardieu *et al*., 2021) and no large-scale, genome-wide studies have been reported to date.

Giant sequoia (*Sequoiadendron giganteum* [Lindl.] J.Buchh.) is a slow-growing, long-lived, outcrossing species that grows in discrete groves on the western slope of the Sierra Nevada mountains in California. Giant sequoia is diploid and has a genome size of 8.125 Gbp (Scott *et al*., 2020). The species occurs in a highly disjunct range consisting of approximately 75 groves, spanning around 420 km north to south and ranging from 830 to 2700 m elevation. Giant sequoia is the most moisture-demanding species of mixed conifer forests, mainly because of its very high leaf area: mature trees can have > 10^8^ leaves (Sillett *et al,.* 2015; Dodd and DeSilva, 2016). Coast redwood (*Sequoia sempervirens* [D.Don] Endl.) is also slow-growing and long-lived but differs from giant sequoia in that it is hexaploid (genome size is 26.5 Gbp; Neale et al. 2021) and often reproduces asexually. The species once had a nearly continuous distribution along the Pacific Coast in Oregon and California, but natural populations were severely reduced by intensive logging beginning in the 19^th^ and 20^th^ centuries (Burns *et al*., 2018; Breidenbach *et al*., 2020). Both coast redwood and giant sequoia are listed as endangered by the International Union for Conservation of Nature (IUCN) Red List of Threatened species (Farjon and Schmid, 2013). However, increased growth rates in response to elevated CO_2_ (Sillett *et al.,* 2015) may make these two species good candidates for forest restoration and carbon sequestration.

Being the tallest and fourth-tallest conifers, the crowns of coast redwood and giant sequoia can stretch over ∼100m of vertical extent, and thus these species have the greatest degree of within-crown phenotypic plasticity of any conifers measured, both responding more strongly to water availability than to light (Chin and Sillett 2016, 2019). Like other members of the Cupressaceae, coast redwood and giant sequoia lack an endodermis to constrain the breadth of their vascular development, allowing the proliferation of traits promoting water-stress tolerance with increasing height (Oldham *et al.,* 2010; Chin and Sillett 2016). Less clear is whether populations of these species have adapted genetically to environmental variation across their ranges, in ways that either limit or enhance phenotypic plasticity in traits related to drought tolerance. In this study, we sampled natural populations across the current ranges of both giant sequoia and coast redwood, grew cuttings in pots in a greenhouse common garden for two years, measured a range of physiological and anatomical traits thought to be relevant for drought resilience, and tested for significant genome-wide associations with ten different drought-related traits using univariate and multivariate GWAS methods. We aimed to dissect the genomic basis of drought tolerance in each species, in order to identify the hardiest individuals and populations that might be used for conservation and restoration efforts in the species.

## RESULTS

### Genotype datasets

A total of 577,774 and 767,242 SNPs were called for 71 SEGI and 82 SESE individuals, respectively. From them, 52,987 (9%) SNPs from 71 SEGI individuals; and 57,357 (7%) SNPs from 82 SESE individuals were retained after filtering using TASSEL. The before and after filtering SNPs statistics for each of the SEGI and SESE individuals are reported in Supplementary Tables S1 and S2, respectively. The filtered SNPs datasets were retained for further GWAS analyses.

**Table 1.** Drought-related traits measured in this study in giant sequoia (SEGI) and coast redwood (SESE).

| Trait | Symbol | Units |
| --- | --- | --- |
| Shoot mass per area | SMA | g m<sup>-2</sup> |
| Osmotic pressure at full turgor | PIFT | MPa |
| C:N ratio | CN | unitless |
| Stable carbon isotope discrimination | D13C | permille |
| Stable nitrogen isotope discrimination | D15N | permille |
| Stomatal density | SD | mm<sup>-2</sup> |
| Total area of transfusion tissue | TA | μm<sup>2</sup> |
| Total area of xylem | XA | μm<sup>2</sup> |
| Total area of central fibers | FA | μm<sup>2</sup> |
| Xylem hydraulic diameter | HD | μm |

### Phenotype datasets

For the phenotypic traits listed in Table 1, within-genotype mean trait values varied widely across genotypes for each species (Figure 1), and showed variation across the species’ natural ranges (Figure 2). In coast redwood, the relative spread of means across genotypes was greatest for central fiber area, C:N ratio and SMA, whereas in giant sequoia the spread was greatest for total areas of transfusion tissue and xylem. In both species, the relative spread was smallest for carbon isotope discrimination and xylem hydraulic diameter. Consistent with established patterns, trait values in giant sequoia indicated a relatively more xeric habit than those in coast redwood; e.g., total xylem and transfusion tissue areas and xylem hydraulic diameter were all smaller in giant sequoia, and osmotic pressure at full turgor and leaf mass per unit area were greater in giant sequoia. Trait variability across genotypes grown in our common garden was generally lower than that seen along vertical gradients within the crowns of individual trees (cf. red bars in Figure 1) (Oldham *et al.,* 2010; Chin and Sillett 2016).

**Figure 1.**
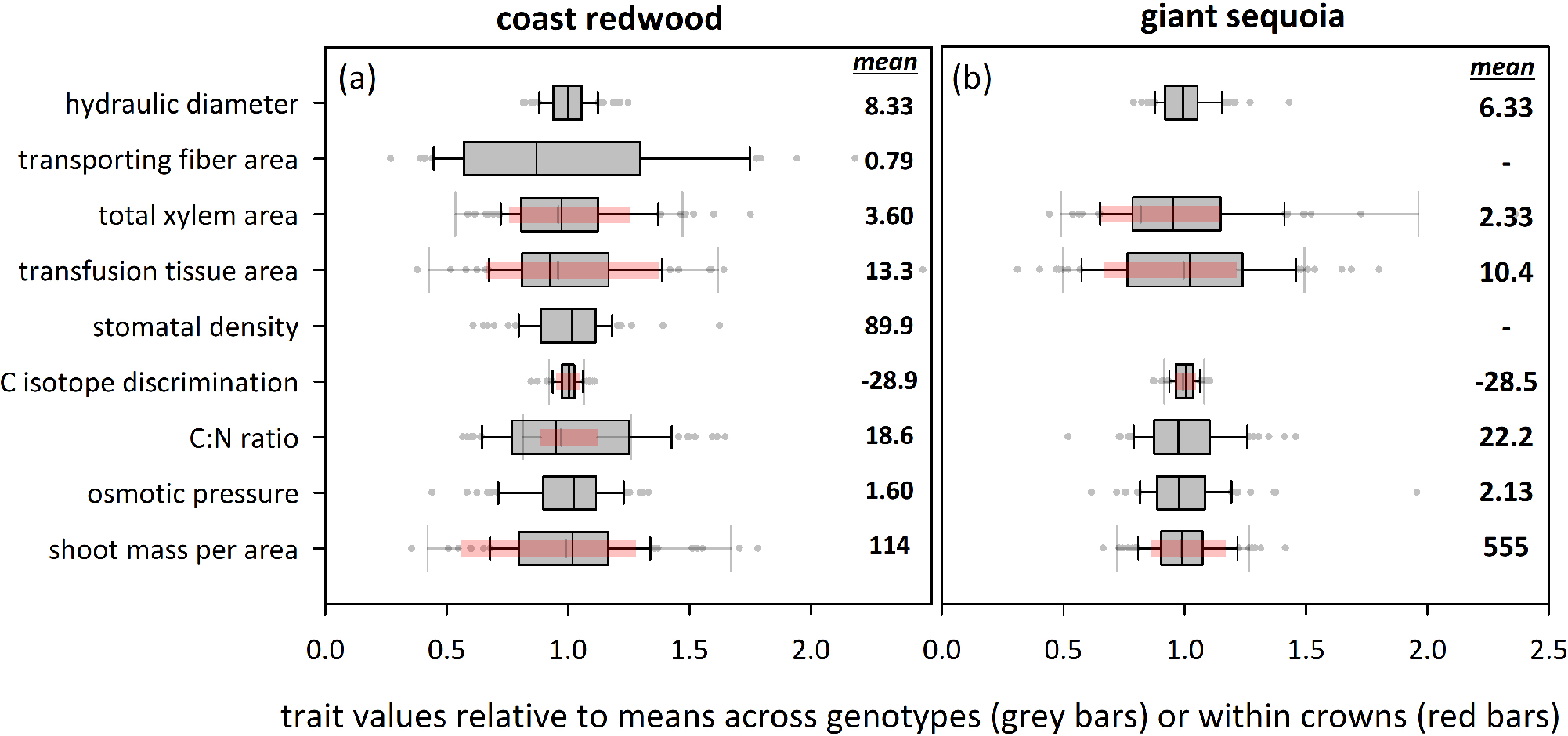
Phenotypic variability in drought-related traits across populations of climatically diverse origin is similar to or smaller than phenotypic plasticity within individual tree crowns. Grey bars indicate variation (interquartile range) across genotypes examined in this study, for (a) coast redwood, and (b) giant sequoia; red bars indicate variation within crowns of single trees examined by Chin & Sillett (2016) and Oldham et al. (2010). The vertical line in each grey bar denotes the median, whiskers denote 5th and 95th percentiles, and grey symbols are outliers. Trait values are expressed relative to mean values across genotypes (grey bars) or within crowns (red bars). Mean values across genotypes for each trait are shown at right in each panel (units as in Table 1; for areas of transporting fibers, xylem and transfusion tissue, multiply values shown here by 10^3^ to get areas in μm^2^).

**Figure 2.**
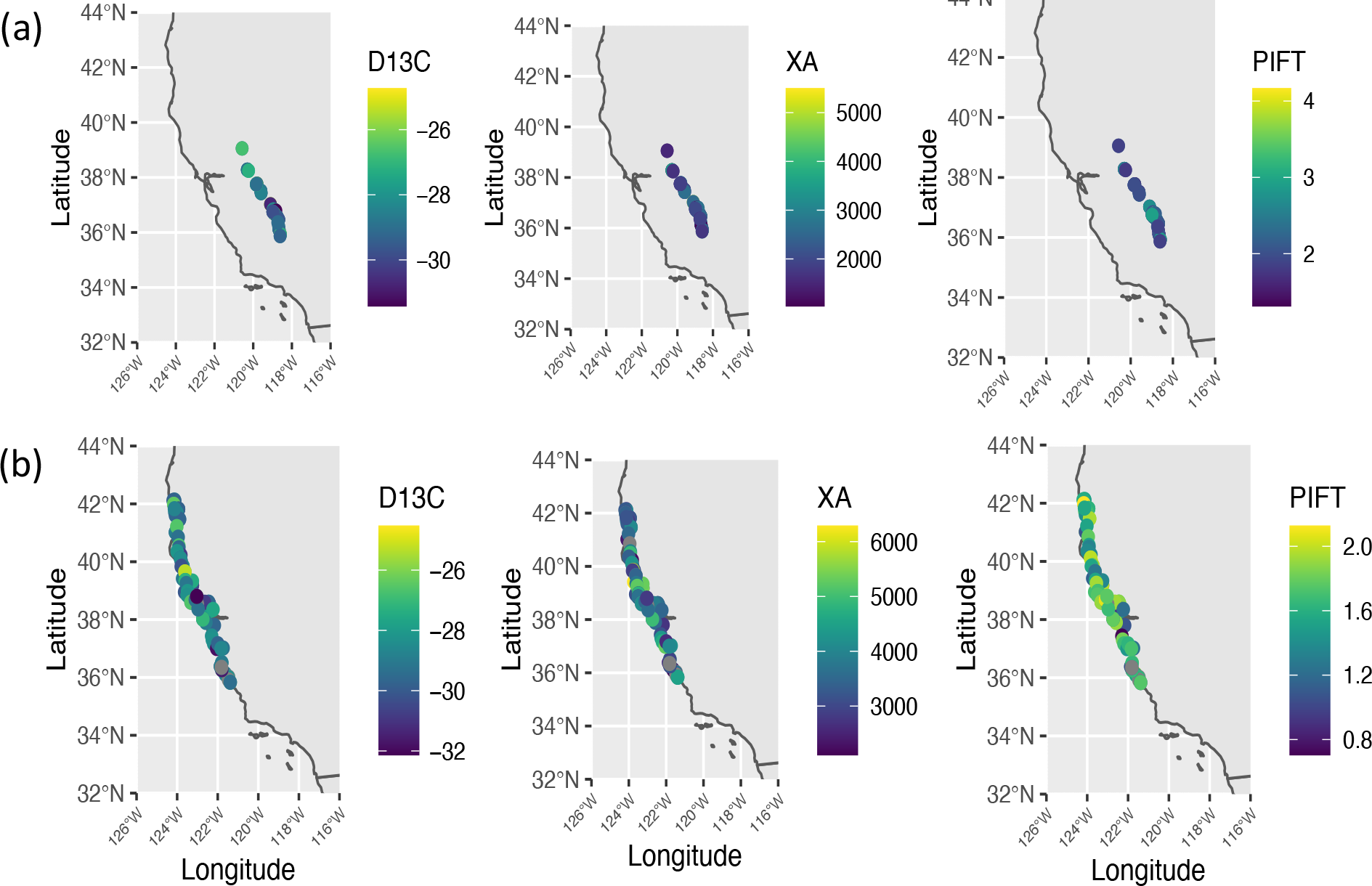
Phenotypic variability across the species’ natural distribution range based on common garden experiments. Carbon isotope discrimination (D13C), total xylem area (XA) and osmotic pressure (PIFT) in (a) giant sequoia and (b) coast redwood.

### Correlations among traits and environmental parameters

In SEGI, C:N ratio (CN) was positively correlated with shoot mass per area (SMA) (r=0.33, p-value=0.004), and carbon isotope discrimination (D13C) (r=0.32, p-value=0.006). Total xylem part of vascular bundle (XA) was also positively correlated with carbon isotope discrimination (r=0.24, p-value=0.04); and total area of transfusion tissue (TA) (r=0.43, p-value=0.0001). All correlations results can be found in Figure 3. Carbon isotope discrimination was positively correlated with latitude (r=0.26, p-value= 0.025) and negatively associated with longitude (r=-0.3, p-value=0.009) and elevation (r=-0.27, p-value=0.021). C:N ratio was negatively correlated with elevation (r=-0.27, p-value=0.019).

**Figure 3.**
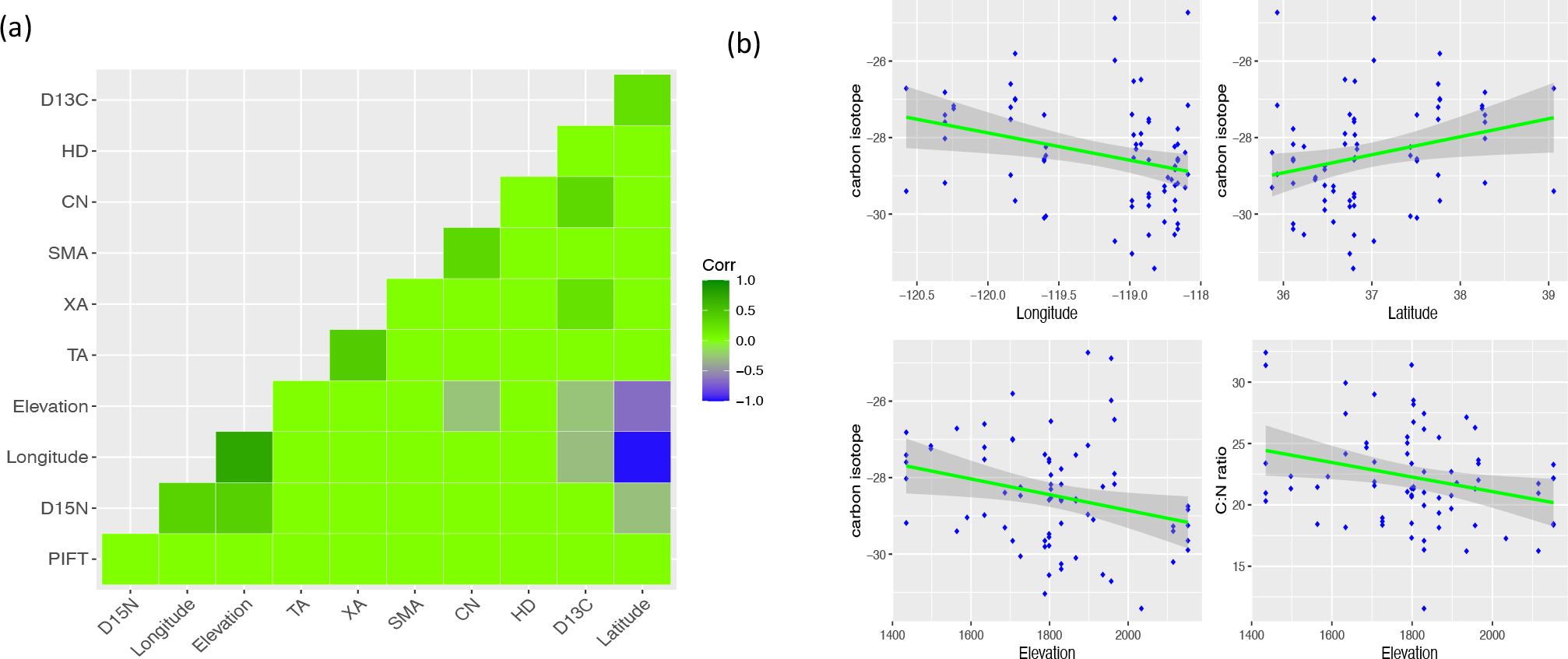
Correlations among drought-related traits and geographical variables in giant sequoia. (a) Heatmap showing R for all combinations of variables; (b) scatterplots of significant correlations (P value<0.05) among geographic variables and carbon isotope discrimination and C:N ratio. Full trait names can be found in Table 1.

Carbon isotope discrimination was positively correlated with several precipitation variables (Bio3, Bio12, Bio13, Bio16, Bio17 and Bio19), suggesting populations at the northern distribution of the species range, located at more humid locations and lower elevations have higher water use efficiency than populations in other locations when grown in a common garden. Osmotic pressure at full turgor (PIFT) was positively correlated with different measures of temperature and precipitation variation (Bio4-Temperature seasonality, Bio7-Temperature Annual Range, Bio15-Precipitation seasonality) and negatively correlated with Relative Humidity (RH). Finally, xylem hydraulic diameter (HD) was correlated with Mean Annual Solar Radiation (MAR). All correlation results can be found in Figure S1.

In SESE, the total area of central fibers (FA) was negatively correlated with latitude (r= -0.3, p-value=0.006), and positively correlated with longitude (r=0.23, p-value=0.037). The same trait was also negatively correlated with various measures of precipitation including MAP, Bio12-Bio19 and positively correlated with several temperature-related variables (MCMT, EMT, Bio3, Bio8, Bio11). Osmotic pressure at full turgor (PIFT) was positively correlated with different precipitation variables such as Bio13, Bio16 and Bio19. Shoot mass per unit area (SMA) was positively correlated with osmotic pressure at full turgor (r = 0.33, p = 0.002), C:N ratio (r = 0.39, p = 0.0003) and total xylem area (r = 0.29, p = 0.01). Finally, xylem hydraulic diameter (HD) was positively correlated with Mean Annual Temperature (MAT), DD5, EMT and Eref, and negatively with degree days below 18 °C (DD_18). All correlation results can be found in Figure S2.

Genotypes from coast redwood’s latitudinal and precipitation extremes had very little overlap within the common-garden trait-space, but in all cases overlapped by at least 70% with intermediate categories; trees originating from intermediate sites thus did not have readily detectable trait differences from either extreme. North and South genotypes shared 19% of their unified trait-space, while Wet and Dry genotypes only intersected in 14% of their unified trait-space, with low levels of overlap indicating multivariate differences in the suites of traits associated with both latitudinal and precipitation extremes. Wet and Dry sites had highly conserved traits compared to Intermediate Precipitation sites, which had 5X less point density within their phenotypic volumes, suggesting a broader hydraulic niche driving much less specialization. Likewise, genotypes from coast redwood’s central latitudes were spread across almost 3X the relative trait-space of North or South genotypes.

### Genome-wide association study (GWAS)

Genome-wide association analyses of 52,987 SNP markers and 71 individuals in SEGI; and 57,357 SNP markers and 82 individuals in SESE were performed to detect marker–trait associations. Mixed linear model (MLM) and general linear mixed model (GLM) were used to determine associations between genotypic and phenotypic datasets in TASSEL. In SEGI, a total number of ∼476K associations were tested among 52.987k SNPs and all phenotypic traits listed in Table 1 (stomatal density [SD] and area of transporting fibers [FA] were excluded from this analysis, in which SD was not measured and transporting fibers were not observed). In SESE, a total of ∼573K associations were tested among 57.357k SNPs and all nine phenotypes. Bonferroni correction for multiple testing was performed to adjust p-values. The general linear mixed model (GLM) identified a total number of 78 significant SNPs, 77 of them were associated with total area of transfusion tissue (TA) and one with total area of transporting fibers (FA) (Table 2, Table S3). These SNPs were distributed across 22 scaffolds and matched 23 genes in the genome of coast redwood (Table S3). In SEGI, GLM only identified 2 SNPs, located at close distance in chromosome 9 and associated with osmotic pressure at full turgor (PIFT) with a p-value < 9.00×10^-6^ after Bonferroni correction (Table 2, Table S4, Figure S3). Manhattan plots of −log10 (P) values for each SNPs versus chromosomal or scaffold positions were generated from these datasets. TASSEL MLM did not identify any significant marker x trait associations in any of the species after Bonferroni correction at threshold p-value < 0.05.

**Table 2.** Functional annotation of the genes of significant SNPs identified by GLM at TASSEL and univariate, multivariate linear mixed models at GEMMA for SESE and SEGI. Analysis column indicates the analysis methods (GLM, uLMM, mvLMM) for finding significant SNPs associated with different phenotypic traits in SEGI and SESE. Similar search ID column indicates the genbank or uniport ID identified by similar BLAST hits. Genes associated with environmental variables in a previous GEA study show the environmental variable in parentheses.

| Analysis | Gene | Similar Search ID<sup>a</sup> | SNPs<sup>b</sup> | p-value | Annotation Description | KEGG Pathway |
| --- | --- | --- | --- | --- | --- | --- |
| GT | SESE_010495<sup>+</sup> | XP_024365529.1 | 1 | 2.00E-07 | F-box protein | Ubiquitin system |
| GT | SESE_022882<sup>+</sup> | XP_024520340.1 | 1 | 3.00E-07 | Uncharacterized protein | Cationic antimicrobial peptide (CAMP) resistance |
| GT | SESE_025289<sup>+</sup> | - | 1 | 6.00E-07 | - | - |
| GT | SESE_026053<sup>+</sup> | XP_006840991.3 | 4 | 3.00E-07 | Motile sperm domain-containing protein | - |
| GU,<br>GM, GT | SESE_026278<sup>+</sup> | XP_006847025.2 | 1 | 1.00E-07 | BTB/POZ domain-containing protein | Hedgehog signaling pathway |
| GT | SESE_028233<sup>+</sup>(MAT) | XP_010255566.1 | 1 | 0.00E+00 | Protein HOTHEAD-like | Glycine, serine and threonine metabolism |
| GT | SESE_031915<sup>+</sup>(MAT) | XP_027156177.1 | 1 | 7.00E-07 | Uncharacterized protein | - |
| GT | SESE_035946<sup>+</sup> | ATG29933.1 | 1 | 1.00E-07 | Cyp750c18 | Cytochrome |
| GU,<br>GM, GT | SESE_039821<sup>+</sup> | XP_012445484.1 | 21 | 2.00E-07 | Receptor-like protein kinase HAIKU2 | - |
| GT | SESE_041334<sup>+</sup> | XP_020251916.1 | 1 | 8.00E-07 | Uncharacterized protein | - |
| GT | SESE_058034<sup>+</sup>(CMD) | XP_011625611.2 | 1 | 0.00E+00 | Cysteine-rich receptor-like protein kinase | - |
| GT | SESE_074679<sup>+</sup> | ABR17724.1 | 1 | 4.00E-07 | Unknown | - |
| GT | SESE_075160<sup>+</sup> | XP_006840304.2 | 1 | 2.00E-07 | Chlorophyll a-b binding protein | Photosynthesis-antenna proteins, Metabolic pathways |
| GT | SESE_079577<sup>+</sup>(MAP) | XP_007014626.2 | 1 | 8.00E-07 | Protein indeterminate domain | - |
| GT | SESE_084919<sup>+</sup> | XP_017700156.2 | 1 | 2.00E-07 | Hypothetical protein | - |
| GT | SESE_093724<sup>+</sup> | XP_006858242.1 | 1 | 1.00E-07 | Glyoxysomal processing protease | - |
| GT | SESE_102359<sup>+</sup> | XP_020532610.1 | 1 | 1.00E-06 | Lysosomal Pro-X carboxypeptidase | Renin-angiotensin system, Protein digestion and absorption |
| GT | SESE_118250<sup>+</sup>(MAP) | XP_027064440.1 | 1 | 4.00E-07 | Uncharacterized protein | - |
| GU,<br>GM, GT | SESE_121791<sup>+</sup> | XP_031476139.1 | 4 | 2.00E-07 | Cysteine protease RD19A-like | Lysosome, Apoptosis, Plant-pathogen interaction |
| GM | SESE_041833 | XP_006838305.1 | 1 | 6.00E-06 | cellulose synthase A catalytic | Transferases |
| GM | SESE_031320<sup>+</sup>(MAP) | XP_027057309.1 | 1 | 4.00E-06 | uncharacterized protein | - |
| GT | SEGI_21288<sup>++</sup>(MAP) | XP_019108059.1 | 1 | 7.18E-07 | Uncharacterized protein | - |
| GM | SEGI_29523 | MBT3334041.1 | 1 | 3.20E-06 | 5-amino-6-(D-ribitylamino) uracil--L-tyrosine 4-hydroxyphenyl transferase-CofH | Methane metabolism, Metabolic pathways |

GU= GEMMA univariate; GM= GEMMA multivariate; GT= GLM at TASSEL; <sup>+</sup>genes were associated with TA; <sup>++</sup> genes were associated with pift phenotype. <sup>a</sup>NCBI or Uniprot IDs; <sup>b</sup>Total Numbers of candidate SNPs in a gene.

**Table 3.** Gene Ontology annotation of the genes of significant SNPs associated with different phenotypic traits in redwood (SESE) and giant sequoia (SEGI).

| Gene | GO IDs | GO Names |
| --- | --- | --- |
| SESE_010495 | MP: GO:0005515 | MP: protein binding |
| SESE_022882 | - | - |
| SESE_026053 | - | - |
| SESE_026278 | MP: GO:0005515 | MP: protein binding |
| SESE_028233 | MP: GO:0016614; MP: GO:0050660 | MP: oxidoreductase activity, acting on CH-OH group of donors; MP: flavin adenine dinucleotide binding |
| SESE_031915 | MP: GO:0003676 | MP: nucleic acid binding |
| SESE_035946 | MP: GO:0004497; MP: GO:0005506; MF: GO:0016705; MP: GO:0020037 | MP: monooxygenase activity; MP: iron ion binding; MP: oxidoreductase activity, acting on paired donors, with incorporation or reduction of molecular oxygen; MP: heme binding |
| SESE_039821 | BP: GO:0006468; MP: GO:0004672; MF: GO:0005524 | BP: protein phosphorylation; MP: protein kinase activity; MP: ATP binding |
| SESE_041334 | CC: GO:0016021 | CC: integral component of membrane |
| SESE_058034 | BP: GO:0006468; MP: GO:0004672; MP: GO:0005524 | BP: protein phosphorylation; MP: protein kinase activity; MP: ATP binding |
| SESE_074679 | CC: GO:0110165 | CC: cellular anatomical entity |
| SESE_075160 | BP: GO:0009765; CC: GO:0016020 | BP: photosynthesis, light harvesting; CC: membrane |
| SESE_079577 | BP: GO:0009630; MP: GO:0046872; CC: GO:0005634 | BP: gravitropism; MP: metal ion binding; CC: nucleus |
| SESE_084919 | - | - |
| SESE_093724 | BP: GO:0016485; MP: GO:0004252; CC: GO:0005777 | BP: protein processing; MP: serine-type endopeptidase activity; CC: peroxisome |
| SESE_102359 | BP: GO:0006508; MP: GO:0008236 | BP: proteolysis; MP: serine-type peptidase activity |
| SESE_118250 | - | - |
| SESE_121791 | BP: GO:0006508; MP: GO:0008234 | BP: proteolysis; MP: cysteine-type peptidase activity |
| SESE_041833 | BP: GO:0030244; MP: GO:0016760; CC: GO:0016020 | BP: cellulose biosynthetic process; MP: cellulose synthase (UDP-forming) activity; CC: membrane |
| SESE_031320 | - | - |
| SEGI_21288 | BP: GO:0006278; BP: GO:0090502; MP: GO:0003676; MP: GO:0003964; MP: GO:0004523 | BP: RNA-dependent DNA biosynthetic process; BP: RNA phosphodiester bond hydrolysis, endonucleolytic; MF: nucleic acid binding; MF: RNA-directed DNA polymerase activity; MF: RNA-DNA hybrid ribonuclease activity |
| SEGI_29523 | MP: GO:0003824; MP: GO:0016740; MP: GO:0016765; MP: GO:0044689; MP: GO:0046872; MP: GO:0051536; MP: GO:0051539 | MP: catalytic activity; MP: transferase activity; MP: transferase activity, transferring alkyl or aryl (other than methyl) groups; MP: 7,8-didemethyl-8-hydroxy-5-deazariboflavin synthase activity; MP: metal ion binding; MP: iron-sulfur cluster binding; MP:4 iron, 4 sulfur cluster binding |

BP=biological process; MP=molecular process; CC=cellular component

Subsequently, univariate linear mixed model (uLMM) and multivariate linear mixed model (mvLMM) approaches were performed in GEMMA to identify significant SNPs. In SESE, mvLMM identified 31 significant SNPs (p-value < 9.00×10^-6^), and uLMM, 29 SNPs (p-value < 9.00×10^-6^) (Figure 4; Tables S5 and S6; Figure S4). Of the 29 SNPs identified from uLMM analysis, 27 were significantly associated with total area of transfusion tissue (TA), one with xylem hydraulic diameter (HD), and one with total area of transporting fibers (FA). In SEGI, mvLMM identified 3 significant SNPs and uLMM only one (p-value < 9.00×10^-6^) associated with total xylem area, XA (Figure 5; Table S7; Figure S5). These SNPs were in chromosomes 5, 8 and 9 of the giant sequoia genome (Table S7). Among all three-analysis including GLM at TASSEL, uLMM and mvLMM at GEMMA, a total 27 significant SNPs were consistently found in SESE (Figure 6). For SEGI, only one significant SNP (chromosome 8) was shared among two of the GWAS analyses (mvLMM and uLMM; Figure 5). Manhattan plots for each SNP versus chromosomal or scaffold positions for GLM (TASSEL), and uLMM and mvLMM (GEMMA) analyses for both species were reported in Figures 4-5, and Figures S3-S5.

**Figure 4.**
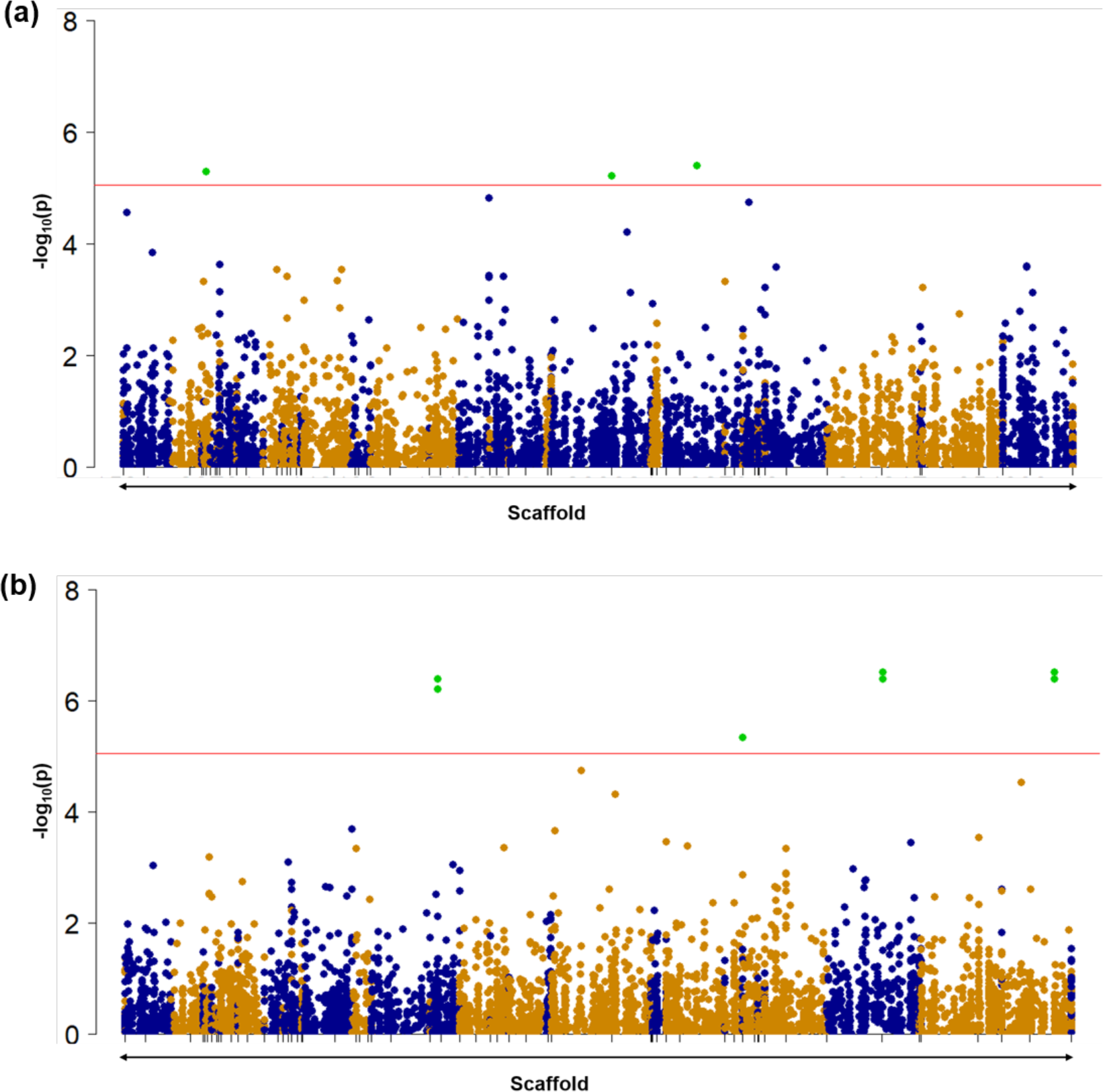
Manhattan plot of SNP markers generated by GEMMA using multivariate linear mixed model (mvLMM) in SESE. (a) manhattan plot indicate the mvLMM analysis with the phenotypic traits SMA, PIFT, SD, D13C, CN and (group 1) (b) manhattan plot indicating the mvLMM analysis with the phenotypic traits D15N, TA, XA, HD and FA (group 2) in SESE. In the Manhattan plot y-axis represent the p-value of SNP markers in -log10 and the x-axis is chromosomal positions. The red line represents genome-wide significant cut-off (p-value < 9.00×10^-6^). The green dot over the genome-wide significant cut-off (red line) represents the significant SNPs (p-value < 9.00×10^-6^).

**Figure 5.**
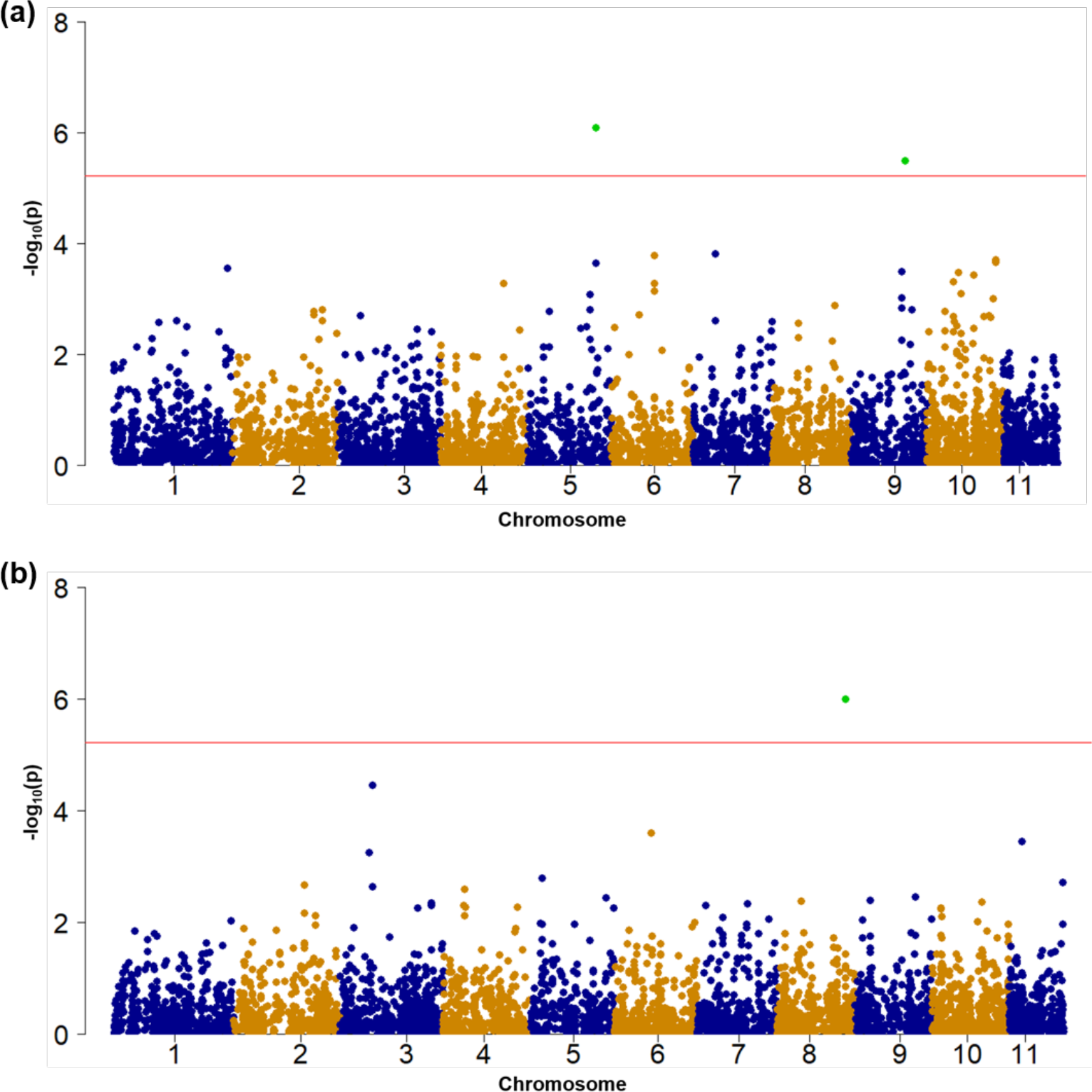
Manhattan plot of SNP markers generated by GEMMA using multivariate linear mixed model (mvLMM) in SEGI. (a) Manhattan plot indicate the mvLMM analysis with the phenotypic traits SMA, PIFT, D13C, CN and D15N (group 1) and (b) manhattan plot indicate the mvLMM analysis with the phenotypic traits TA, XA, HD (group 2) in SEGI. The red line represents genome-wide significant cut-off (p-value< 6.00×10^-7^). The green dot over the genome-wide significant cut-off (red line) represents the significant SNPs (p-value< 6.00×10^-7^).

**Figure 6.**
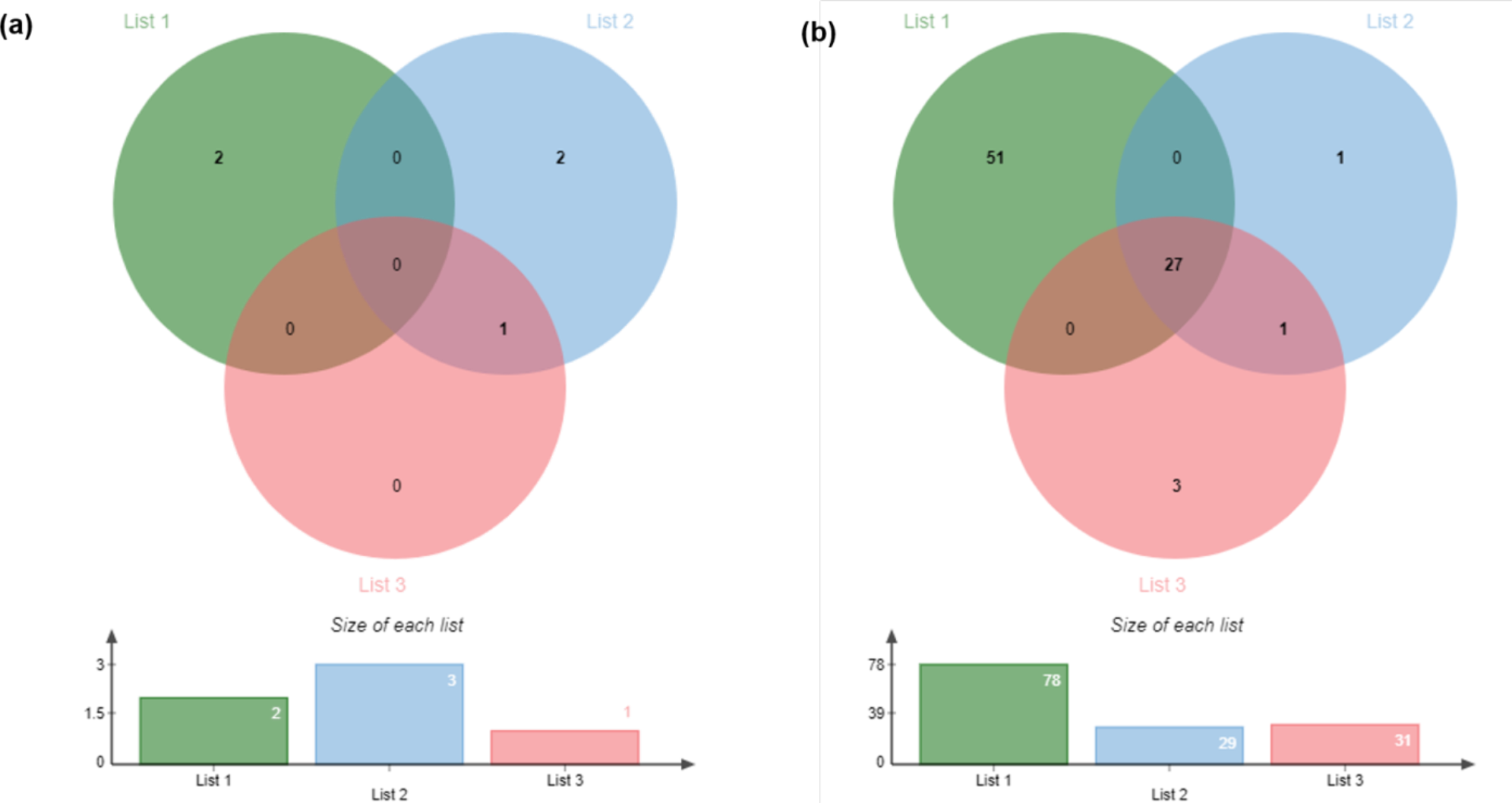
**Venn-diagrams** representing the common significant SNPs identified by all three GWAS analyses including GLM at TASSEL, uLMM and mvLMM at GEMMA in SEGI (a) and SESE (b). List 1 is the total number of SNPs identified by GLM, list 2 is total number of SNPs identified by uLMM and list 3 is total SNPs identified by mvLMM.

In SESE, all significant SNPs associated with TA, came from genes involved in the ubiquitin system, cationic antimicrobial peptide (CAMP) resistance, hedgehog signaling pathway, glycine, serine and threonine metabolism, lysosome, apoptosis, plant-pathogen interaction, renin-angiotensin system, and protein digestion and absorption (Table 3). In SESE, gene SESE_010495 was annotated as a F-box protein, SESE_026053 as a motile sperm domain-containing protein, SESE_026278 as a BTB/POZ domain-containing protein, SESE_028233 as a protein HOTHEAD-like, and SESE_039821 as a receptor-like protein kinase HAIKU2 (Table 3). In SEGI, a significant SNP at the gene SEGI_21288 associated with PIFT was identified as an uncharacterized protein. The GO IDs and GO names of these genes of significant SNPs in SESE and SEGI were reported in Table 3.

## DISCUSSION

By using a combination of univariate and multivariate methods, our study was able to identify several genes associated with drought-related traits in two ecologically important conifer species: coast redwood and giant sequoia. Previous genome-wide studies identifying candidate genes for drought tolerance have been absent for both of these important species. Here, we report notable phenotypic variation for several drought-related traits among natural populations (or groves) of both giant sequoia and coast redwood grown in a common garden. The development of genome-wide methods, gene identification, functional annotation and location in the species’ genomes has only been possible due to the recent sequencing of both species’ reference genomes (Scott *et al*., 2020; Neale *et al*., 2021).

### Polygenic basis of drought tolerance

Our results suggest a polygenic basis of drought tolerance, consistent with previous genome-wide association studies in other complex traits in conifer species, with candidate genes distributed in different chromosomes or scaffolds, and small to moderate effect sizes (Baison *et al*., 2020; Weiss *et al*., 2020; De La Torre *et al*., 2021a). The exact location of candidate genes in the genome of coast redwood could only be determined at the scaffold level since the current assembly of the reference genome is not chromosome-scale (Neale *et al*., 2021). Univariate methods identified a total of 78 new significant associations for coast redwood, 27 of them were consistently found by all three univariate and multivariate GWAS methods in this study. All these SNPs (except one) were associated with variations in the total area of transfusion tissue associated with the leaf vasculature. However, when using the multi-trait multivariate mvLMM method in GEMMA, we found that many of the SNPs identified by TASSEL were associated with a group of drought-related traits, either vascular or carbon isotope-related traits. This is coincident with the presence of significant correlations among traits in these groups (Figure 3), suggesting the multitrait multivariate GWAS provides a more accurate picture of the complex trait architecture in the species. Six genes associated with total area of transfusion tissue in coast redwood were also associated with Mean Annual Temperature, Mean Annual Precipitation or Climate Moisture Deficit in a previous environmental genome-wide association study using the same SNP set (Table 2; De la Torre *et al*., 2021b). Transfusion tissue area triples with height in tall coast redwood crowns, and is associated with low water availability and less-negative values of δ^13^C; it buckles under drought stress, releasing water that helps protect the leaf and isolate damage (Oldham *et al*.,. 2010).

The number of significant associations was much lower in giant sequoia with only six significant associations discovered by all GWAS methods, with two of them associated with osmotic pressure at full turgor, one with the total xylem area, and three with combinations of traits (Figure 5). The multitrait multivariate mvLMM method did not result in significant differences or a higher number of candidate genes when compared with the univariate methods. Significant genes were involved in RNA-dependent DNA biosynthetic processes and catalytic activity (Table 3). One of the genes, SEGI_21288 (chromosome 9) associated with osmotic pressure, was also associated with Mean Annual Precipitation in a previous GEA study in giant sequoia (De La Torre *et al*., 2021b).

In coast redwood, most associations were clustered in a small number of scaffolds and genes. For example, scaffold 16773 harbors three closely located genes (SESE_102359, SESE_121791, SESE_010570) involved in proteolysis (Table 3); scaffold 203021 has four different genes (SESE_026053, SESE_008114, SESE_041334 and SESE_031915) with unknown functions, and scaffold 344217 has two genes (SESE_039821 and SESE_025289), the first one, a receptor-like protein kinase HAIKU2 involved in protein phosphorylation, and the second one with unknown function. The identification of other potential genomic clusters could only be possible with the presence of a chromosome-scale genome assembly in coast redwood. No genomic clusters were observed in giant sequoia mainly due to the small number of significant associations.

Despite the relatively small sample size of the common garden experiments, substantial phenotypic variation was found in several of the drought-related traits measured in this study. For example, C:N ratio, total area of transfusion tissue, total xylem area, and total area of central conducting fibers showed great variation in the species (Figure 1). This large variation, however, did not translate into the identification of large numbers of candidate genes. There might be several explanations for this: the presence of high levels of plasticity for these drought-related traits in the species; a large difference between the number of markers and samples leading to false positives after stringent multiple testing correction; or the relatively low power to detect rare variants due to small sample sizes. Due to the genome-wide distribution and the number of markers included in this study, we don’t consider the number of markers to be a potential limitation in our study despite the rapid decay of linkage disequilibrium in the species.

### High levels of phenotypic variance in drought-related traits

Giant sequoia is known for its high phenotypic plasticity and multiple adaptations to cope with water stress, including shoot and leaf succulence, leaf toughness, tight stomatal control of water loss and increasing xylem cavitation resistance with height (Pitterman *et al*., 2012; Ambrose *et al*., 2015; Chin and Sillet 2016). In a greenhouse study, Ambrose *et al*., (2016) found contrasting drought-responses strategies between the species, with greater stomatal closure leading to an increase in intrinsic water-use efficiency and lower xylem embolism under severe drought in giant sequoia than in coast redwood. As an adaptation to their natural environment, shade-tolerant coast redwood seedlings will invest biomass into above-ground woody stem, which enhances competitive success in humid, closed canopy conditions with shallow water tables seen in northern forests (Sawyer *et al*., 2000; Ambrose *et al*., 2016). In contrast, though larger as adults, giant sequoia seedlings invest more biomass in developing root growth as desiccation is an important factor contributing to early mortality in the species (Havey *et al*., 1980).

A study measuring shoot water potential, leaf gas exchange, xylem embolism and growth concluded there were not significant differences at the population level in neither coast redwood nor giant sequoia (Ambrose *et al*., 2016). In contrast, our study found significant population-level differences in three traits for each species (carbon isotope discrimination, osmotic pressure, and xylem hydraulic diameter for giant sequoia, and total area of central fibers, osmotic pressure and xylem hydraulic diameter for coast redwood). For example, carbon isotope discrimination of bulk leaf tissue for plants grown in our common garden was positively correlated with several precipitation and geographic variables for each genotype’s environment of origin. For a given photosynthetic capacity, a decrease in carbon isotope discrimination (i.e., less negative values of D13C) implies reduced stomatal opening and greater water use efficiency. Thus, our results suggest that giant sequoia genotypes collected from sites near the species’ northern limit, or from more humid or lower-elevation (< 2000 m) sites, have higher water use efficiency when grown in a common garden than genotypes from more southern, drier or higher-elevation sites. Under these criteria, we identified three high-elevation groves that might need conservation due to a higher sensitivity to drought (given lower carbon isotope values and lower water use efficiency) should their year-round supplies of surface water diminish; these are Redwood Mountain (36.69 latitude, -118.92 longitude), Giant Forest (36.56 latitude, -118.75 longitude) and Atwell Mill (36.46 latitude, -118.68 longitude). All these groves are located over 2000 m of elevation, where cold tolerance traits, such as narrow xylem tracheid diameters, may have been selected for over those supporting drought survival.

Within-crown phenotypic plasticity in tall trees of both coast redwood and giant sequoia is only slightly greater than that observed in the common garden, for comparable traits (red bars in Figure 1; Oldham *et al*., 2010; Chin and Sillett 2016). The innate ability to acclimate to environmental microclimatic conditions, and the mostly small differences in within-crown compared to within-garden variation suggests that naturally recruiting trees of northern provenance may have a different range of plasticity less suited to withstand climatic pressures comparable to conditions experienced in the southern range. Indeed, southernmost coast redwood trees reach a maximum height 20-30 m shorter than northern trees but have similar treetop levels of transfusion tissue investment (Ishii *et al*., 2014). In contrast to the two species explored here, Douglas-fir (*Pseudotsuga menziesii*) has varying amounts of within-crown trait variability across its much larger range (Chin and Sillett 2019), suggesting that future genomic work may find tradeoffs between geographic and individual-level trait variation.

In coast redwood, larger values for two traits associated with the capacity for water transport – the total area of conducting fibers contributing to water transport, and the xylem hydraulic diameter (a measure of the effective mean size of individual xylem conduits, accounting for nonlinear effects of conduit size on water transport) – were associated with lower precipitation and higher temperatures in the environment of origin for genotypes. For example, genotypes collected from lower latitudes and more eastern locations (stands at Warm Springs Creek [38.68 latitude, -123.11 longitude] and Bodega [38.36 latitude, -122.96 longitude]) had particularly large areas devoted to central conducting fibers. Central fibers are found in coast redwood at their greatest abundance in a distinct shoot morphotype, specialized for absorption of water (Alana Chin, unpublished data), so the increased area at dry sites may indicate a reliance on summertime foliar water uptake and use of alternate hydraulic pathways. These traits may indicate adaptations that enable water transport to be sustained in environments that are relatively warm and dry for this species; sustained water transport would, in turn, minimize leaf water stress and enable leaf stomata to remain open to allow photosynthesis (Brodribb *et al*., 2007). Thus, these groves might represent sources of drought-tolerant germplasm for coast redwood.

Coast redwood genotypes from wet and dry locations have distinct combinations of water-stress related functional traits when considered on a multivariate level, and far less variability than seen among intermediate rainfall sites – suggesting adaptive specialization. Intraspecific trait convergence is a characteristic response to abiotic stress and so is expected on environmentally harsh, typically dry or cold, range ends (Mitchell and Baaker 2014, Van Nulan *et al*., 2020). In the case of coast redwood, rainforest conditions may present unique challenges due to months of continuous leaf wetness and heavy cloud cover, resulting in phenotypes on both latitudinal range-ends that overlap with intermediate zones, but share little of the same trait-space. Better group separation based on precipitation-class, rather than latitude, may indicate that climatic adaptation has been more important than distance between populations in determining the coast redwood water-stress phenotype. Latitudinal groupings may be undetectable in giant sequoia because of the relatively consistent climate within its range; coast redwood samples came from sites spanning >2.5X the climatic variability (based on mean coefficient of variation).

### Functional annotation of candidate genes

Candidate genes found in this study indicate a complex genomic architecture of drought tolerance with many genes involved in many important biological functions related to growth, abiotic stress resistance, and disease resistance. For example, gene SESE_010495 associated with total area of transfusion tissue was involved in the ubiquitin system. The ubiquitin-proteasome system controls the degradation of most proteins in the cells. It provides a rapid strategy to control many cellular processes by degrading specific proteins, playing a critical role in the regulation of many biological processes such as hormonal signaling, growth, embryogenesis, senescence, and environmental stress (Sharma *et al*., 2016; Xu and Xue 2019). F-box domain proteins have been found to play important roles in abiotic stress responses via the ubiquitin pathway. For example, the study by Zhou *et al*., (2015) found that overexpression of *TaFBA1* enhanced drought tolerance in transgenic plants, confirming the importance of F-box proteins in plants’ tolerance to multiple stress conditions.

A significant SNP at the gene SESE_026278 which annotated as BTB/POZ domain-containing protein was involved in hedgehog signaling pathway. The BTB/POZ domain is an evolutionarily conserved and widely distributed structural motif found involved in different biological processes, such as transcriptional regulation, cytoskeletal organization, and formation of voltage-gated channels (Collins *et al*., 2001). Overexpression of GmBTB/POZ in soybean resulted in enhanced resistance to *Phytophthora sojae* (*P. sojae*). The activities and expression levels of enzymatic superoxide dismutase (SOD) and peroxidase (POD) antioxidants were significantly higher in GmBTB/POZ-overexpressing transgenic soybean than in wild-type (WT) plants (Zhang *et al*., 2019).

Another important candidate gene identified in our study was SESE_039821, a receptor-like protein kinase. Receptor-like kinases are important signaling components that regulate a variety of cellular processes. Protein kinases regulate metabolic pathways and are intimately involved in cellular signaling networks (Wang *et al.,* 2007). An Arabidopsis cDNA microarray analysis led to the identification of the cysteine-rich receptor-like kinase CRK36 responsive to the necrotrophic fungal pathogen (*Alternaria brassicicola*) (Lee *et al.,* 2017). The gene *haiku2* is a mutant allele of gene *iku2*, which is a leucine-rich repeat kinase (LRR) gene involving in regulation of seed size in Arabidopsis (Luo *et al*., 2005).

In our study, the gene SESE_121791 was identified as cysteine protease RD19A-like and was involved in lysosome, apoptosis and plant-pathogen interaction pathways. Papain-like cysteine proteases (PLCPs) are involved in many plant processes (Zou *et al*., 2018). Cysteine proteases were found to play a role in nodule development in soybean and in the pathogen defense (Shukla *et al*., 2014; van Wyk *et al*., 2014). Also, cysteine protease (*AdCP*) gene in the wild peanut (*Arachis diogoi*) was differentially expressed when it was challenged with the late leaf spot pathogen (Shukla *et al*., 2014).

The gene SESE_075160 identified by GLM at TASSEL in our study was identified as chlorophyll a/b binding protein. The light-harvesting chlorophyll *a*/*b*-binding (LHCB) members were shown to be targets of an ABA-responsive WRKY-domain transcription factor, which represses *LHCB* expression to balance the positive function of the LHCBs in ABA signaling. Consequently, it revealed that ABA is an inducer that fine-tunes *LHCB* expression through repressing the WRKY40 transcription repressor in stressful conditions in co-operation with light, which allows plants to adapt to environmental challenges (Liu *et al*., 2013).

This study is a step forward to understand the genomics of drought tolerance in long-generation conifer species. Genomic studies have been limited in conifers due to their large genome sizes, and long-generation times. Given the high levels of phenotypic variance despite the relatively small sample sizes in both coast redwood and giant sequoia found in this study, long-term studies with larger sample sizes are warranted. For that purpose, coast redwood seedlings measured in this study have been planted in long-term common gardens in California, where different phenotypes can be evaluated as trees mature. This new resource, together with our newly sequenced reference genomes of giant sequoia and coast redwood (Scott *et al*., 2020; Neale *et al*., 2021) will help develop future genomic studies in the species. A thorough knowledge of the interconnection among plasticity, genomics and physiological processes is needed to predict species’ responses to future warmer conditions and to design conservation and management strategies.

## EXPERIMENTAL PROCEDURES

### Foliage collection for greenhouse establishment

Juvenile foliage of coast redwood (SESE) was collected from the Kuser common garden (Kuser *et al*., 1995) hedge orchard growing in Russell Reserve (University of California field station, Contra Costa County, California) in fall 2017. As the SESE-Kuser common garden is hedged annually, juvenile primary shoots were collected for ideal propagation. Cuttings were taken of foliage from mature giant sequoia (SEGI) trees in the Fins trial (Fins, 1979) at Foresthill Divide Seed Orchard (Foresthill, California) in winter 2018. As the SEGI accessions were mature trees, juvenile foliage was sampled where possible, but sampling was restricted to plagiotropic growth. Collections were made to represent a wide range of geographic sites of origin, spanning the species’ natural distributions.

Immediately following collection, foliage samples were misted with water, wrapped in paper towels, and stored in labeled plastic bags. Bagged samples were then kept in a cooler with ice for transport to the greenhouse, where they were stored in a refrigerated room (4 °C) for up to 24 hours. One at a time, to avoid mixing of genotypes, samples were washed with water to remove debris, then briefly soaked in a disinfectant (Physan 20, solution of 39 mL L^-1^). Terminal shoots were then trimmed into cuttings approximately 10 cm long. All primary needles were removed from the lower third of the each cutting. Between 30 and 60 cuttings per genotype were stuck. Cuttings were dipped in rooting hormone (3:1 Dip N Grow:water, 7,500 PPM IBA) for 5 seconds and then stuck into rooting medium (9:1 perlite:peat by volume) with Osmocote 18-6-12 controlled release fertilizer at 1.8 kg m^-3^ and Micromax Micronutrients at 0.7 kg m^-3^. Cuttings were arranged in rows, with 3 cm between individual cuttings, and a minimum of 5 cm between rows. Rooting trays were kept under mist until roots emerged (for SESE, 2-3 months; for SEGI, 4 months or longer). Rooted cuttings were carefully removed from rooting medium and potted into individual containers with growing medium, individually labeled, staked with bamboo if needed, and returned to the greenhouse. Re-potted plants were hand watered for 3-4 weeks and then placed on irrigation drip.

For SESE genotyping, fresh needles were collected from a selected ramet from each of the surviving 92 clonal genets. Overall, these samples were sourced from 66 locations, with 1-3 source trees per population. For SEGI genotyping, fresh needles were sampled from a selected ramet from each of the surviving 90 clonal genets. These SEGI accessions came from 23 groves, with 1-9 samples from each population. Additionally, six SEGI accessions were included as technical replicates, resulting in 96 genotyped samples total.

### DNA extraction

Young needles were collected from a selected ramet from each of the surviving 92 SESE and 90 SEGI genets. They were stored on ice for transport, then flash-frozen in liquid nitrogen, stored in a -80°C freezer for 48 hours, and lyophilized (48 hours for SEGI & 72 hours for SESE). Global DNA (gDNA) was extracted with the Omega Biotek E-Z 96 Plant DNA kit and an Eppendorf automated pipetting workstation at UC Davis. The DNA extraction protocol included one day of tissue lysis, followed by several steps of precipitation, filtering and elution. DNA quality was assessed using a Qubit 2.0 Fluorometer (average concentration = 24.5 ng/μL for SESE and 43.5 ng/μL for SEGI), NanoDrop 8000 (average A260/280 = 1.94; average A260/230 = 1.99 for SESE and 1.6 for SEGI), and gel electrophoresis (average fragment size ≥ 20,000 bp). Samples were normalized to 20 ng/μL in 50 μL. The gDNA was submitted to the UC Davis Genome Center for sonication, size selection, and library preparation.

### Sequence Capture and SNP calling

Exome capture baits were designed for each species using PacBio IsoSeq RNA data combined with previously published Illumina RNAseq data (Scott *et al*., 2020) and clustered at 95% identity to produce a set of nonredundant transcripts. The clustered transcripts were then mapped to the reference assembly at high stringency using gmap. For SEGI, the regions of matches were submitted to Roche (Madison, WI) where 120-mer oligos were designed to cover the target regions at 2x tiling density. For SESE, the regions of matches between genome sequence and transcript sequences were submitted to Roche for 120-mer oligos were designed to cover the target regions at 2x tiling density. The UC Davis Genome center carried out hybridization of baits and the gDNA samples described above. The resulting libraries were pooled and sequenced on the NovaSeq 6000 platform. Bowtie2 v2.2.9 (Langmead and Salzberg, 2012) was used to align sequencing capture raw reads against the reference genome assemblies of giant sequoia version 2.0 (treegenes.db.org/FTP/Genomes/Segi) and coast redwood version 2.1 (treegenesdb.org/FTP/Genomes/Sese). Alignments were sorted and divided into multiple sets based on reference intervals, and later processed in parallel using SAMtools v1.3.1 and BEDtools v2.25.0. SNPs were then called using BCFtools with default parameters (Li et al., 2009). Haplotypes were called using Genome Analysis Toolkit (GATK v.4.1.7.0) HaplotypeCaller & GenotypeGVCF (McKenna et al., 2010). SNP functional annotations were obtained from the species’ reference genome annotations in the TreeGenes database (treegenesdb.org); and by sequence alignment against the NCBI non-redundant protein sequences database (nr) using BLASTP (Johnson et al., 2008) with an e-value <1x10^-10^. BCFtools was used to merge vcfs files of individuals for further analysis (Danecek et al., 2011).

### Phenotypic traits

For each species, we measured ten traits related to drought tolerance (Table 1) in one branch from each of three individuals per genotype. The set of available individuals from each genotype were distributed randomly throughout the greenhouse; sampling was performed haphazardly, in that we sampled the first three individuals encountered for each genotype. In some cases, this required exhaustive searching due to poor survival of some genotypes; in other genotypes, many individuals were present. Each branch was sampled in early June 2020 using sharp secateurs and immediately placed in a ziploc bag and sprayed with water. The bag was then sealed and placed in a cooler with ice to prevent further water loss. Upon return to the laboratory, each branch was recut under water (≥ 1.5 cm), the cut end was placed into a 50-mL falcon tube, and the tube was placed into a stand to allow the branch to rehydrate for 38 – 46 hours. After rehydration, three leaves were removed from each branch and stored in FAA for later anatomical measurements, and the branch was immediately returned to a sealed ziploc bag that had been sprayed internally with water. These bags were stored in a refrigerator until completion of measurement of shoot mass per unit area and subsampling for osmotic pressure measurements and stomatal density mounts were completed. Three values (or, in a few cases, two) for each trait measurement were thus collected for each genotype, and subsequent analysis was performed on the mean of these three values. Methods for each trait measurement are described below.

### Shoot mass per unit area

Each branch was removed from the refrigerator and its sealed bag, and a small, representative section was returned to the bag and refrigerator for subsequent measurements of osmotic pressure and stomatal density. The rest of the branch was dabbed dry with paper towels and placed on a scanner (Canon TR8520), scanned for later measurement of shoot silhouette area (including both leaves and the shoots to which they were attached) in ImageJ, placed into a labelled paper envelope, and placed in a drying oven at 70 °C until weight stopped declining (generally ∼24 h). These dried samples were later weighed on a 5-point digital balance (Mettler-Toledo model XS225DU). Shoot mass per unit area was computed as the ratio of dry mass to initial (fresh) silhouette area.

### Stomatal density

For SESE, three leaves from each branch were excised and mounted abaxial side down in fingernail polish on a microscope slide. The number of stomata in a single image frame at a magnification of 200x was counted for each leaf and divided by the frame size (0.255742 mm^2^) to calculate stomatal density. Results are presented as the mean ± SE among leaves. Stomatal density was not measured for SEGI.

### Osmotic pressure at full turgor

For each branch, a 6-mm long section of a previously rehydrated leaf (SESE) or branch (SEGI) was excised with a fresh razor blade and immediately enclosed in the sample well of a C-52 thermocouple psychrometer (Wescor, Logan, UT). The psychrometer was then placed in an insulated box and allowed to equilibrate. Every hour, a CR6 datalogger (Campbell Scientific, Logan, UT) was used to initiate a 10-s cooling curve for each psychrometer, psychrometer output (μV) was recorded every second, and the average μV output between 2 and 5 seconds after the end of cooling was calculated. The resulting means were found to remain stable between 4 and 9 hours of equilibration; values from either 5 or 6 hours were used for subsequent analysis. Each psychrometer was calibrated using five KCl solutions, with osmotic pressures of 0, 0.5, 1.0, 2.0, and 3.0 MPa, with 0.025 mL of each solution placed on a filter paper disk in the psychrometer sample well and otherwise measured as described earlier for leaves.

### Elemental and isotopic analyses

A dried sample of leaf (SESE) or branch (SEGI) material was placed in a sealed cuvette with three stainless steel spherical pellets and ground in a ball mill for two minutes. Subsamples (1.9 – 4.6 mg) were weighed and transferred into tin capsules, placed into 96-well trays, and crushed to seal the capsules. δ^13^C (relative to Vienna Pee Dee Belemnite standard) and total C and N were measured at the UC Davis Stable Isotope Facility using a PDZ Europa ANCA-GSL elemental analyzer interfaced to a PDZ Europa 20-20 isotope ratio mass spectrometer (Sercon Ltd., Cheshire, UK), with several replicates of at least four laboratory reference standards periodically interspersed for internal calibration. C:N ratio (mol mol^-1^) was calculated by dividing total C by total N.

### Leaf vascular anatomy

Leaves previously stored in FAA as described earlier were hand-sectioned, mounted on a slide, and digitally imaged at 400x magnification, centered on the single leaf vein, and four traits were measured using ImageJ: (i) the total cross-sectional area of transfusion tissue laterally abutting the single leaf vein, (ii) the total cross-sectional area occupied by xylem, (iii) the hydraulic mean diameter (calculated following Kolb and Sperry 1999 as HD = Σ*D*^5^/Σ*D*^4^, where *D* is conduit diameter and the sum is taken over 10 conduits; *D* was calculated from conduit lumen area [*A*] as *D* = [4*A*/*π*]^0.5^), and (iv) the total area of central fibers, when present (longitudinal fibers with thick, concentrically lamellated cell walls located adjacent to the adaxial side of the xylem, and are thought to contribute to water transport). Central fibers were not observed in SEGI. As for all other traits, measurements were repeated for three leaves per genotype, each taken from a different ramet.

### Correlations among drought-related traits, geographic and environmental variables

Physiological parameters depend on relationships among traits and their composite effects on leaf function; thus, we evaluated geo-climatic clustering within the collective phenotypic trait-space observed in the common garden. We also tested and plotted correlations among the nine drought-related traits (Table 1) and geographical and environmental variables for both SEGI and SESE using R packages Hmisc and ggcorrplot in R studio 1.1.442. Geographic variables (latitude, longitude and elevation) representing the geographic origin of the sampled trees (collected directly for SESE individuals, or using the centroid of the grove polygon for SEGI) were used as geographic origin to obtain environmental data from public databases such as WorldClim2.0 (Fick and Hijmans, 2017) and ClimateNA (Wang *et al*., 2016).

All 83 SESE genotypes were ordinated in Euclidean trait-space with PCA, using a correlation matrix and eight of nine traits, excluding only C:N because of univariate nonlinear relationships with other traits. A similar analysis was repeated unsuccessfully for SEGI, which did not have any geographic or climatic associations with PCA axes. Giant sequoia had only five traits suitable for PCA, giving less dimensionality to explore, and samples came from far fewer groves which were sampled unevenly. Grove-level clusters were apparent in the trait-space, but our sample size did not permit analysis on that level. The eight SESE traits used had a mean skewness of 0.366 and a mean kurtosis of 0.596. None of the climatic or geographic variables had strong correlations with individual axes; however, the cumulative association of rainfall-related variables and latitude were > R^2^ = 0.3. We selected mean annual precipitation and latitude to create two sets of potential clusters within the trait-space, selecting three groups from each, with the highest and lowest values forming two groups, and the intermediate values forming a larger, third class. For latitude we called genotypes from above 40° latitude “north” (*N* = 23), those from latitudes below 37.5° “south” (*N* = 19), and intermediate zones “central” (*N* = 41). Categories for rainfall were “wet” if sampling sites received more than 1600 mm of annual precipitation (*N* = 19), those from locations with fewer than 900 mm of precipitation “dry” (*N* = 16), and “intermediate rainfall” (*N* = 48). With the first five PCA axes, retaining 78% of the total trait variation, we found the 5-dimensional “phenotypic volumes” as minimum convex hulls occupied by each potential latitudinal or rainfall class, and estimated their intersection and union using the R package hypervolume.

### Genotype data preparation

Raw genotyping data containing high levels of missing data were filtered using TASSEL v.5.2.72 (Bradbury *et al*., 2007) with the following parameters: minor allele frequency (maf) = 0.05, maximum allele frequency (max-maf) = 0.9. The minimum count–the minimum number of taxa in which the site must have been scored to be included in the filtered data set, 50 was implemented for SEGI and 30 for SESE.

### Genome-wide association study

Associations between each drought-related trait and individual marker were tested using a general linear model (GLM) and a mixed linear model (MLM) implemented in the GWAS analysis in TASSEL v.5.2.72 (Bradbury *et al*., 2007). A kinship matrix and Principal Component Analysis (PCA) were calculated for the mixed linear model (MLM) analysis (Yu *et al*., 2006). Population structure was accounted by including principal components as co-variates in the models. Relatedness among individuals was also accounted for incorporating a kinship matrix in the models. Effect sizes (proportion of phenotypic variance explained by the marker) and the dominance and additive effects were also calculated in TASSEL.

In addition, univariate linear mixed models (uLMM) and multivariate linear mixed models (mvLMM) GWAS were performed in GEMMA v0.98.3 (Zhou and Stephens 2012; 2014). In contrast to the uLMM method, mvLMM associates multiple phenotypic traits with all markers simultaneously, while controlling for population structure and relatedness. To run GEMMA, PLINK binary ped format was generated using PLINK v.1.9 software for association analysis. The Bonferroni threshold (<0.05) correction and false discovery rate (FDR) were applied for multiple correction to identify significant SNPs. Manhattan plots of −log10 (P) values for each SNPs versus chromosomal positions were generated at the GLM of TASSEL and uLMM and mvLMM of GEMMA results.

### Functional gene annotations

The genomic positions of the significant SNPs were investigated to identify the annotated genes by scanning the genomic VCF files of SEGI and SESE. Subsequently, the identified significant SNPs were annotated using annotation files downloaded from TreeGenes (https://treegenesdb.org/TripalContactProfile/588450). The annotation was confirmed using some other approaches such as pfam (Finn *et al*., 2014) and blastp (Johnson *et al*., 2008), BlastKOALA (Kanehisa *et al*., 2016). The Pfam was ran using the HMMER (Finn *et al*., 2011) at default parameters with e-value 1.0 to search proteins families. The blastp was ran at expected threshold-0.05; matrix-BLOSUM 62; database- non-redundant protein sequence (nr) to search the similar hits. The BlastKOALA at KEGG (Kanehisa *et al*., 2016) was performed for protein pathways and annotations. The identical matching genes were chosen to identify annotations and KEGG pathways.

## DATA AVAILABILITY STATEMENT

Sequencing raw reads are deposited in the NCBI SRA (https://www.ncbi.nlm.nih.gov.sra) under BioSample SUB10142549.

## ACKNOWLEDGEMENTS

The authors would like to thank Emily Burns for her guidance and expertise on early stages of the project, Bill Libby for help on collection design, and Bill Werner for help with greenhouse support and propagation. TNB thanks Zane Moore for providing measurements of stomatal density in SESE, and Oliver Betz for producing micrographs of leaf cross sections in SESE and SEGI. This project was supported by a grant from Save The Redwoods League for the Redwood Genome Project (to DN). ARDLT was supported by NIFA grant ARZZ19-0258. TNB was supported by the National Science foundation (Awards #1557906 and 1951244) and the USDA National Institute of Food and Agriculture (Hatch project 1016439 and Award no. 2020-67013-30913).

## AUTHOR CONTRIBUTIONS

DN, ARDLT, TNB and AS designed the study; AS established the common gardens; TNB, AS, and AROC planned the trait measurement; TNB measured all drought-related traits; BA and AS performed all genomic lab work; DP and SS did the SNP calling; MKS and ARDLT performed all genomic and bioinformatic data analyses; MKS, TNB, AROC and ARDLT wrote the manuscript; all authors reviewed the final version of the manuscript.

## SUPPORTING INFORMATION

**Table S1.** SNP filtering in giant sequoia

**Table S2.** SNP filtering in coast redwood.

**Table S3.** Univariate GLM TASSEL GWAS results in coast redwood.

**Table S4.** Univariate GLM TASSEL GWAS results in giant sequoia.

**Table S5.** Univariate uLMM GEMMA GWAS results in coast redwood.

**Table S6.** Multitrait multivariate mvLMM GEMMA GWAS results in coast redwood.

**Table S7.** uLMM and mvLMM GEMMA results in giant sequoia.

**Figure S1.** Correlations among drought related traits, geographic and environmental variables in giant sequoia. R values are color-coded based on the figure legend.

**Figure S2.** Correlations among drought related traits, geographic and environmental variables in coast redwood. R values are color-coded based on the figure legend.

**Figure S3.** Manhattan plot of SNP markers generated by TASSEL using GLM for SEGI (a) for SESE (b). Each point represents a genetic variant in Manhattan plots. In the Manhattan plot y-axis represent the p-value of SNP markers in -log10 and the x-axis is chromosomal positions.

**Figure S4.** Manhattan plot of SNP markers generated by GEMMA using univariate linear mixed model (uLMM) in SESE. (a) manhattan plot indicate the uLMM analysis with the phenotypic traits CN, (b) D13C, (c) D15N, (d) FA, (e) SMA, (f) PIFT, (g) SD, (h) TA, (i) HD, and (j) XA, in SESE.

In the Manhattan plot y-axis represents the p-value of SNP markers in -log10 and the x-axis is chromosomal positions. The red line represents genome-wide significant cut-off (p-value < 9.00E10^-6^). The green dot over the genome-wide significant cut-off (red line) represents the significant SNPs (p-value < 9.00E10^-6^).

**Figure S5.** Manhattan plot of SNP markers generated by GEMMA using univariate linear mixed model (uLMM) in SEGI. (a-h) Manhattan plot indicate the uLMM analysis with the phenotypic traits (a) TA, (b) CN, (c) D13C, (d) D15N, (e) SMA, (f) PIFT, (g) HD, and (h) XA, in SEGI. In the Manhattan plot y-axis represent the p-value of SNP markers in -log10 and the x-axis is chromosomal positions. The red line represents genome-wide significant cut-off (p-value< 6.00E10^-7^). The green dot over the genome-wide significant cut-off (red line) represents the significant SNPs (p-value< 6.00E10^-7^).

## REFERENCES

1. Adams, H. D., Zeppel, M. J., Anderegg, W. R., Hartmann, H., Landhäusser, S. M., Tissue, D. T., et al (2017) A multi-species synthesis of physiological mechanisms in drought-induced tree mortality. *Nat*. Ecol. Evol, 1(9), 1285–1291.

2. Adams, H.D., and Kolb, T.E. (2005) Tree growth response to drought and temperature along an elevation gradient on a mountain landscape. J. Biogeogr., 32, 1629–1640.

3. Allen, C.D., Macalady, A.K., Chenchouni, H., Bachelet, D., McDowell, N., Vennetier, M., Kitzberger, T., Rigling, A., Breshears, D.D., Hogg, E.T. Gonzalez, P. (2010) A global overview of drought and heat-induced tree mortality reveals emerging climate change risks for forests. For. Ecol. Manag, 259(4), 660–684.

4. Ambrose, A.R., Baxter, W.L., Wong, C.S., Burgess, S.S.O., Williams, C.B., Næsborg, R.R., Koch, G.W., Dawson, T.E. (2016) Hydraulic constraints modify optimal photosynthetic profiles in giant sequoia trees. Oecologia, 182, 713–730

5. Ambrose, A.R., Baxter, W.L., Wong, C.S., Næsborg, R.R., Williams, C.B., Dawson T.E. (2015) Contrasting drought-response strategies in California redwoods. Tree Physiol, 35, 453–469

6. Baison, J., Zhou, L., Forsberg, N. et al. (2020). Genetic control of tracheid properties in Norway spruce wood. Sci Rep, 10, 18089. https://doi.org/10.1038/s41598-020-72586-3

7. Bellard, C., Bertelsmeier, C., Leadley, P., Thuiller, W., and Courchamp, F. (2012). Impacts of climate change on the future of biodiversity. Ecol Lett, 15, 365–377.

8. Bradbury, P. J., Zhang, Z., Kroon, D.E., Casstevens, T.M., Ramdoss, Y., Buckler, E.S. (2007) TASSEL: software for association mapping of complex traits in diverse samples. Bioinformatics, 23 (19), 2633–5, doi: 10.1093/bioinformatics/btm308.

9. Breidenbach N, Gailing O, Krutovsky KV (2020) Genetic structure of coast redwood (*Sequoia sempervirens* [D. Don] Endl.) populations in and outside of the natural distribution range based on nuclear and chloroplast microsatellite markers. PLOS ONE, 15(12), e0243556. https://doi.org/10.1371/journal.pone.0243556

10. Brodribb, T.J., Field, T.S., Jordan, G.J (2007) Leaf maximum photosynthetic rate and venation are linked by hydraulics. Plant Physiol, 144, 1890–1898

11. Burns, E. E., Campbell, R., and Cowan, P. D. (2018). State of Redwoods Conservation Report: A Tale of Two Forests, Coast Redwoods, Giant Sequoia. Save the Redwoods League, San Francisco CA USA.

12. Chen, ZQ., Zan, Y., Milesi, P. et al. Leveraging breeding programs and genomic data in Norway spruce (*Picea abies* L. Karst) for GWAS analysis. Genome Biol 22, 179 (2021). https://doi.org/10.1186/s13059-021-02392-1

13. Chin, A.R. and Sillett, S.C. (2016) Phenotypic plasticity of leaves enhances water-stress tolerance and promotes hydraulic conductivity in a tall conifer. Am. J. Bot., 103, 796–807

14. Chin, A.R. and Sillett, S.C. (2019). Within-crown plasticity in leaf traits among the tallest conifers. Am. J. Bot., 106(2), 174–186.

15. Collins, T., Stone, J. R., Williams, A.J (2001). All in the family: the BTB/POZ, KRAB, and SCAN domains. Mol. Cell. Biol., 21 (11), 3609–3615, doi:10.1128/Mcb.21.11.3609-3615.2001.

16. Cumbie, W. P., Eckert, A., Wegrzyn, J., Whetten, R., Neale, D., and Goldfarb, B. (2011). Association genetics of carbon isotope discrimination, height and foliar nitrogen in a natural population of *Pinus taeda* L. Heredity, 107, 105–114.

17. Danecek, P., Auton, A., Abecasis, G., Albers, C.A., Banks, E., DePristo, M.A., Handsaker, R.E., Lunter, G., Marth, G.T., Sherry, S.T., McVean, G., Durbin, R and 1000 Genomes Project Anal Grp. (2011) The variant call format and VCFtools. Bioinformatics, 27 (15), 2156-2158, doi: 10.1093/bioinformatics/btr330.

18. De La Torre, A.R., Birol, I., Bousquet, J., Ingvarsson, P.K., Jansson, S., Jones, S.J.M., Keeling, C.I., MacKay, J., Nilsson, O., Ritland, K., Street, N., Yanchuk, A., Zerbe, P., Bohlmann, J. (2014) Insights into Conifer Giga-genomes. Plant Physiol, 166, 1–9.

19. De La Torre, A.R., Wilhite, B., Puiu, D., St. Clair, J.B., Crepeau, M.W., Salzberg, S.L., Langley, C.H., Allen, B., Neale, D.B. (2021a) Dissecting the polygenic basis of cold adaptation using genome-wide association of traits and environmental data in Douglas-fir. Genes, 12:110, doi:10.3390/genes12010110.

20. De La Torre, A.R., Sekhwal, M.K., Neale, D.B. (2021b). Selective sweeps and polygenic adaptation drive local adaptation along moisture and temperature gradients in natural populations of coast redwood and giant sequoia. **Genes**. In review

21. Depardieu, C., Gérardi, S., Nadeau, S., Parent, G.J., Mackay, J., Lenz, P., Lamothe, M., Girardin, M.P., Bousquet, J. and Isabel, N. (2021), Connecting tree-ring phenotypes, genetic associations and transcriptomics to decipher the genomic architecture of drought adaptation in a widespread conifer. Mol Ecol, 30: 3898–3917. https://doi.org/10.1111/mec.15846

22. Dodd, R.S. and DeSilva, R. (2016) Long-term demographic decline and late glacial divergence in a Californian paleoendemic: *Sequoiadendron giganteum* (giant sequoia). Ecol Evol, 6, 3342–3355. https://doi.org/10.1002/ece3.2122

23. Eckert, A.J., van Heerwaarden, J., Wegrzyn, J.L., Nelson, C.D., Ross-Ibarra, J., González-Martínez, S.C., Neale, D.B. (2010) Patterns of Population Structure and Environmental Associations to Aridity Across the Range of Loblolly Pine (*Pinus taeda* L., Pinaceae), Genetics, Volume 185, Issue 3, 1 July 2010, Pages 969–982, https://doi.org/10.1534/genetics.110.115543

24. Elfstrand, M., Baison, J., Lundén, K., et al. (2020) Association genetics identifies a specifically regulated Norway spruce laccase gene, *PaLAC5*, linked to *Heterobasidion parviporum* resistance. Plant Cell Environ, 43, 1779–1791. https://doi.org/10.1111/pce.13768

25. Farjon, A., and Schmid, R. (2013). Sequoia sempervirens. The IUCN Red List of Threatened Species., e.T34051A2841558.

26. Fettig, C. J., Mortenson, L. A., Bulaon, B. M., Foulk, P. B. (2019). Tree mortality following drought in the central and southern Sierra Nevada, California, US. For Ecol Manag, 432, 164–178.

27. Fick, S.E. and Hijmans, R.J. (2017), WorldClim 2: new 1-km spatial resolution climate surfaces for global land areas. Int. J. Climatol, 37, 4302–4315. https://doi.org/10.1002/joc.5086

28. Finn, R. D., Bateman, A., Clements, J., Coggill, P., Eberhardt, R.Y., Eddy, S.R., Heger, A., Hetherington, K., Holm, L., Mistry, J., Sonnhammer, E.L., Tate, J., Punta, M. (2014) Pfam: the protein families database. Nucleic Acids Res, 42 (Database issue), D222–30, doi: 10.1093/nar/gkt1223.

29. Finn, R. D., Clements, J., Eddy, S.R. (2011) HMMER web server: interactive sequence similarity searching.” Nucleic Acids Res, 39 (Web Server issue), W29-37, doi: 10.1093/nar/gkr367.

30. Gaylord, M.L., Kolb, T.E., Pockman, W.T., Plaut, J.A., Yepez, E.A., Macalady, A.K., Pangle, R.E., McDowell, N.G. (2013) Drought predisposes piñon-juniper woodlands to insect attacks and mortality. New Phytol, 198(2), 567–578.

31. Gonzalez-Martinez, S. C., Huber, D., Ersoz, E., Davis, J. M., Neale, D. B. (2008). Association genetics in Pinus taeda L. II. Carbon isotope discrimination. Heredity, 101, 19–26.

32. Kolb, K. J. and Sperry, J. S. (1999). Differences in drought adaptation between subspecies of sagebrush (*Artemisia tridentata*). Ecology, 80(7), 2373–2384.

33. Kuser, J.E., Bailly, A., Franclet, A., Libby, W.J., Rydelius, J., Schoenike, R., Vagle, N (1995) Early results of a range wide provenance test of *Sequoia sempervirens*. For Genet Resour, 23, 21–25

34. Harvey, H.T., Shellhammer, H.S., Stecker, R.E. (1980) Giant sequoia ecology. Science Monograph Series 12. US Department of the Interior, National Park Service, Washington, DC.

35. Hicke, J.A., Meddens, A.J.H., Kolden, C.A. (2015) Recent tree mortality in the western United States from bark Beetles and forest fires. For Scie, 62(2),141–153

36. Ishii, H.R., Azuma, W., Kuroda, K. and Sillett, S.C. (2014) Pushing the limits to tree height: could foliar water storage compensate for hydraulic constraints in *Sequoia sempervirens*?. Funct. Ecol, 28(5), 1087–1093.

37. Jactel, H., Petit, J., Desprez-Loustau, M., Delzon, S., Piou, D., Battisti, A., Koricheva, J. (2012) Drought effects on damage by forest insects and pathogens: a meta-analysis. Glob. Change Biol, 18(1), 267–276.

38. Johnson, M., Zaretskaya, I., Raytselis, Y., Merezhuk, Y., McGinnis, S., Madden, T.L. (2008) NCBI BLAST: a better web interface. Nucleic Acids Res, 36 (Web Server issue), W5-9, doi: 10.1093/nar/gkn201.

39. Kanehisa, M., Sato, Y., Kawashima, M., Furumichi, M., Tanabe, M. (2016) KEGG as a reference resource for gene and protein annotation. Nucleic Acids Res, 44 (D1), D457–D462, doi: 10.1093/nar/gkv1070.

40. Kanehisa, M., Sato, Y., Morishima, K. (2016). BlastKOALA and GhostKOALA: KEGG Tools for Functional Characterization of Genome and Metagenome Sequences. J Mol Biol, 428, (4):726–731, doi: 10.1016/j.jmb.2015.11.006.

41. Langmead, B. and S. L. Salzberg. (2012) Fast gapped-read alignment with Bowtie 2. Nat Methods, 9 (4), 357–9, doi: 10.1038/nmeth.1923.

42. Lee, D. S., Kim, Y.C., Kwon, C.J., Ryu, C.M., Park, O.K. (2017) The Arabidopsis Cysteine-Rich Receptor-Like Kinase CRK36 Regulates Immunity through Interaction with the Cytoplasmic Kinase BIK1. Front Plant Sci, 8, doi: ARTN 1856

43. Li, H., Handsaker, B., Wysoker, A., Fennell, T., Ruan, J., Homer, N., Marth, G., Abecasis, G., Durbin, R and Subgroup Genome Project Data Processing. (2009) The Sequence Alignment/Map format and SAMtools. Bioinformatics, 25, (16), 2078-9, doi: 10.1093/bioinformatics/btp352.

44. Liu, R., Xu, H.R., Jiang, S.C., Lu, K., Lu, Y.F., Feng, X.J., Wu, Z., Liang, S., Yu, Y.T., Wang, X.F., Zhang, D.P. (2013) Light-harvesting chlorophyll a/b-binding proteins, positively involved in abscisic acid signalling, require a transcription repressor, WRKY40, to balance their function. J. Exp. Bot., 64 (18), 5443-5456, doi: 10.1093/jxb/ert307.

45. Lu, M., Krutovsky, K.V., Nelson, C.D., West, J.B., Reilly, N.A., Loopstra, C.A. (2017) Association genetics of growth and adaptive traits in loblolly pine (*Pinus tae*da L.) using whole-exome-discovered polymorphisms. Tree Genet, 13, 57.

46. Luo, M., Dennis, E.S., Berger, F., Peacock, W.J., Chaudhury, A. (2005) MINISEED3 (MINI3), a WRKY family gene, and HAIKU2 (IKU2), a leucine-rich repeat (LRR) KINASE gene, are regulators of seed size in Arabidopsis. Proc Natl Acad Sci U S A, 102 (48), 17531–6, doi: 10.1073/pnas.0508418102.

47. McCarthy, M. I., Abecasis, G. R., Cardon, L. R., Goldstein, D. B., Little, J., Ioannidis, J. P., and Hirschhorn, J. N. (2008). Genome-wide association studies for complex traits: consensus, uncertainty and challenges. Nat Rev Genet, 9, 356–69.

48. McGuire, A.L., Gabriel, S., Tishkoff, S.A. et al. (2020) The road ahead in genetics and genomics. Nat. Rev. Genet. 21, 581–596.

49. McKenna, A., Hanna, M., Banks, E., Sivachenko, A., Cibulskis, K., Kernytsky, A., Garimella, K., Altshuler, D., Gabriel, S., Daly, M., DePristo, M.A. (2010). The genome analysis toolkit: a MapReduce framework for analyzing next-generation DNA sequencing data. Genome Res, 20,1297–1303.

50. Mitchell, R.M. and Bakker, J.D. (2014) Intraspecific trait variation driven by plasticity and ontogeny in *Hypochaeris radi*cata. PloS one, 9(10), p.e109870.

51. Moran, E., Lauder, J., Musser, C., Stathos, A., Shu, M. (2017) The genetics of drought tolerance in conifers. New Phytol, 216(4), 1034–1048.

52. Murray, B. G., Leitch, I. J., Bennett, M. D. (2004). Gymnosperm DNA C-values database.

53. Myles, S., Peiffer, J., Brown, P.J., Ersoz, E.S., Zhang, Z., Costich, D.E. and Buckler, E.S. (2009) Association mapping: critical considerations shift from genotyping to experimental design. Plant Cell. 21, 2194-2202.

54. Neale, D.B., Zimin, A.V., Zaman, S., Scott, A.D., Shrestha, B., Workman, R.E., Puiu, D., Allen, B.J., Sekhwal, M.K., De La Torre, A.R., McGuire, P.E., Burns, E., Timp, W., Wegrzyn, J.L., Salzberg, S.L. (2021). Assembled and annotated 26.5 Gbp coast redwood genome: a resource for estimating evolutionary adaptive potential and investigating hexaploidy origin. G3: Genes Genomes Genetics. In press.

55. Neale, D. B. and Savolainen, O. (2004). Association genetics of complex traits in conifers. Trends Plant Sci, 9, 325–30.

56. Oldham, A.R., Sillett, S.C., Tomescu, A.M., Koch, G.W. (2010) The hydrostatic gradient, not light availability, drives height-related variation in *Sequoia sempervirens* (Cupressaceae) leaf anatomy. Am. J. Bot, 97(7), 1087–1097.

57. Pittermann, J., Stuart, S.A., Dawson, T.E., Moreau, A. (2012) Cenozoic climate change shaped the evolutionary ecophysiology of the Cupressaceae conifers. Proc Natl Acad Sci USA 109: 9647–9652.

58. Pritchard, J.K., Stephens, M., Rosenberg, N.A. and Donnelly, P. (2000) Association mapping in structured populations. Am. J. Hum. Genet. 67, 170–181.

59. Quinlan, A. R. and Hall, I.M. (2010) BEDTools: a flexible suite of utilities for comparing genomic features. Bioinformatics, 26 (6), 841–2, doi: 10.1093/bioinformatics/btq033.

60. Razgour, O., Forester, B., Taggart, J. B., Bekaert, M., Juste, J., Ibanez, C., Puechmaille, S. J., Novella-Fernandez, R., Alberdi, A., and Manel, S. (2019). Considering adaptive genetic variation in climate change vulnerability assessment reduces species range loss projections. Proc Natl Acad Sci USA, 116, 10418–10423.

61. Sawyer JO; Sillett SC; Libby WJ et al. (2000) Redwood trees, communities, and ecosystems: a closer look. In: Noss RF (ed) The redwood forest: history, ecology, and conservation of the coast redwoods. Island Press, Washington, DC, 81–118.

62. Scott, A.D., Zimin, A.V., Puiu, D., Workman, R., Britton, M., Zaman, S., Caballero, M., Read, A.C., Bogdanove, A.J., Burns, E., Wegrzyn, J., Timp, W., Salzberg, S.L., Neale, D.B. (2020) Reference Genome Sequence for Giant Sequoia, G3 GENES GENOM GENET,10 (11), 3907–3919, https://doi.org/10.1534/g3.120.401612

63. Sharma, B., Joshi, D., Yadav, P.K., Gupta, A.K., Bhatt, T.K. (2016) Role of Ubiquitin-Mediated Degradation System in Plant Biology. Front Plant Sci, 7:806, doi: 10.3389/fpls.2016.00806.

64. Shukla, P., Singh, N.K., Kumar, D., Vijayan, S., Ahmed, I., Kirti, P.B. (2014) Expression of a pathogen-induced cysteine protease (AdCP) in tapetum results in male sterility in transgenic tobacco. Funct Integr Genomics, 14 (2), 307–17, doi: 10.1007/s10142-014-0367-2.

65. Sillett, S.C., Van Pelt, R., Carroll, A.L., Kramer, R.D., Ambrose, A.R., Trask, D.A. (2015) How do tree structure and old age affect growth potential of California redwoods?. Ecol. Monogr., 85(2), 181–212.

66. Stephenson, N. L., Das, A. J., Ampersee, N. J., Cahill, K. G., Caprio, A. C., Sanders, J. E., Williams, A. P. (2018). Patterns and correlates of giant sequoia foliage dieback during California’s 2012-2016 hotter drought. For. Ecol. Manag, 419, 268–278.

67. Trujillo-Moya, C., George, J.P., Fluch, S., Geburek, T., Grabner, M., Karanitsch-Ackerl, S., Konrad, H., Mayer, K., Sehr, E.M., Wischnitzki, E., Schueler, S. (2018) Drought Sensitivity of Norway Spruce at the Species’ Warmest Fringe: Quantitative and Molecular Analysis Reveals High Genetic Variation Among and Within Provenances. G3 GENES GENOM GENET, 8 (4), 1225–1245, https://doi.org/10.1534/g3.117.300524

68. Van Nuland, M.E., Vincent, J.B., Ware, I.M., Mueller, L.O., Bayliss, S.L., Beals, K.K., Schweitzer, J.A., Bailey, J.K. (2020) Intraspecific trait variation across elevation predicts a widespread tree species’ climate niche and range limits. Ecol. Evol, 10(9), pp.3856–3867.

69. Van Wyk, S. G., Du Plessis, M., Cullis, C.A., Kunert, K.J., Vorster. B.J. (2014) Cysteine protease and cystatin expression and activity during soybean nodule development and senescence. BMC Plant Biol, 14:294, doi: 10.1186/s12870-014-0294-3.

70. Wang, H., Chevalier, D., Larue, C., Cho, S.K., Walker, J.C. (2007) The Protein Phosphatases and Protein Kinases of *Arabidopsis thaliana*. Arabidopsis Book no. 5, e0106, doi: 10.1199/tab.0106.

71. Wang T, Hamann A, Spittlehouse D, Carroll C (2016) Locally Downscaled and Spatially Customizable Climate Data for Historical and Future Periods for North America. PLOS ONE 11(6): e0156720. https://doi.org/10.1371/journal.pone.0156720

72. Weiss, M., Sniezko, R., Puiu, D., Crepeau, M.W., Stevens, K., Salzberg, S.L., Langley, C.H., Neale, D.B., De La Torre, A.R. (2020) Genomic basis of white pine blister rust quantitative disease resistance and its relationship with qualitative resistance. Plant J, doi: 10.1111/tpj.14928.

73. Xu, F. Q. and H. W. Xue. (2019) The ubiquitin-proteasome system in plant responses to environments. Plant Cell Environ, 42 (10), 2931–2944, doi: 10.1111/pce.13633.

74. Yu, J. M., Pressoir, G., Briggs, W.H., Bi, I.V., Yamasaki, M., Doebley, J.F., McMullen, M.D., et al (2006) A unified mixed-model method for association mapping that accounts for multiple levels of relatedness. Nat. Genet., 38 (2), 203–208, doi: 10.1038/ng1702.

75. Zhang, C., Gao, H., Li, R., Han, D., Wang, L., Wu, J., Xu, P., Zhang, S. (2019) GmBTB/POZ, a novel BTB/POZ domain-containing nuclear protein, positively regulates the response of soybean to Phytophthora sojae infection. Mol Plant Pathol, 20 (1), 78–91, doi: 10.1111/mpp.12741.

76. Zhou, S. M., Kong, X.Z., Kang, H.H., Sun, X.D., Wang, W. (2015) The involvement of wheat F-box protein gene TaFBA1 in the oxidative stress tolerance of plants. PLoS One, 10 (4), e0122117, doi: 10.1371/journal.pone.0122117.

77. Zhou, X. and M. Stephens. (2012) Genome-wide efficient mixed-model analysis for association studies. Nat. Genet, 44, 821–824.

78. Zhou, X. and M. Stephens. (2014) Efficient multivariate linear mixed model algorithms for genome-wide association studies. Nat. Methods, 11, 407–409.

79. Zou, Z., Huang, Q., Xie, G., Yang, L. (2018) Genome-wide comparative analysis of papain-like cysteine protease family genes in castor bean and physic nut. Sci Rep, 8 (1), 331, doi: 10.1038/s41598-017-18760-6.

